# HPInet: Interpretable prediction of Host-Pathogen protein-protein Interactions using a transformer-based neural network

**DOI:** 10.1101/2025.08.02.668281

**Authors:** Yueming Hu, Liya Liu, Yanyan Zhu, Enyan Liu, Haoyu Chao, Sida Li, Cong Feng, Yanshi Hu, Yuhao Chen, Shilong Zhang, Yifan Chen, Luyao Xie, Yejun Wang, Ming Chen

## Abstract

Gram-negative bacteria utilize a series of secretion systems (T1SS-T10SS) to deliver secreted effector proteins (T1SE-T10SE) into host cells, leading to infections and diseases. Understanding the interactions between these effector proteins and host proteins is crucial for unraveling the pathogenic mechanisms of bacterial pathogens. Despite advancements in sequencing technologies that have significantly enhanced our knowledge of effector protein diversity and structure, the molecular mechanisms underlying their interactions with human host proteins remain unclear. Experimental approaches to validate these interactions are labor-intensive, time-consuming, and insufficient to comprehensively map the extensive effector-host protein interaction (PPI) network. Additionally, specialized computational tools for predicting these interactions remain scarce. To address this gap, we developed HPInet, a transformer-based deep learning model for predicting interactions between bacterial effector proteins and human host proteins. HPInet introduces convolutional layers and global response normalization (GRN) into the feed-forward network (FFN) of the transformer architecture, enhancing the extraction of local sequence features. Compared to existing state-of-the-art methods, HPInet significantly improves prediction performance, increasing accuracy from 0.502 to 0.891 and boosting sensitivity by 35.3%. Furthermore, by leveraging the model’s attention mechanism, HPInet identifies critical residues at interaction sites, demonstrating its capability to capture local structural features of PPI sites solely from sequence information. To facilitate its application in research, HPInet has been implemented as a freely accessible web server available at https://bis.zju.edu.cn/hpinet. This platform provides a powerful and interpretable tool for studying effector-host protein interactions, offering valuable insights into bacterial pathogenesis and potential therapeutic targets.

## Introduction

Infectious diseases remain a major global health threat, causing millions of deaths annually. These diseases are often closely linked to the ability of pathogens to invade and manipulate host cells [1]. Understanding the mechanisms of pathogen-host interactions is crucial for uncovering infection patterns and pathogenic mechanisms [2]. Pathogens achieve this by interacting with host proteins, disrupting physiological functions, and triggering pathological responses. Therefore, elucidating and predicting pathogen-host protein-protein interactions (PPIs) is essential for understanding pathogenesis and identifying novel therapeutic targets [2].

Gram-negative bacteria, an important group of human pathogens, utilize secretion systems (T1SS-T10SS) to deliver effector proteins (T1SE-T10SE) into host cells [3–5]. These effector proteins interact with host proteins through diverse mechanisms, manipulating signaling pathways, immune responses, and metabolic functions. For instance, *Salmonella* effector proteins SopE and SopE2, secreted via the type III secretion system (T3SS), interact with host RhoGTPases to induce cytoskeletal remodeling and trigger enteritis [6]. Similarly, *Yersinia* effector YopJ inhibits host immune responses and induces cell death through acetylation of target proteins, contributing to plague pathology [7, 8]. *Helicobacter pylori* effector CagA, delivered via the type IV secretion system (T4SS), disrupts host signaling pathways and promotes gastric cancer development [9]. These studies underscore the critical role of bacterial effectors in pathogenesis and highlight the need to understand their interactions with host proteins.

Despite their importance, experimental validation of bacterial effector-host PPIs remains a significant challenge. Although techniques such as yeast two-hybrid (Y2H), co-immunoprecipitation (Co-IP), and mass spectrometry (MS) have been developed to detect PPIs [10], these methods are labor-intensive, time-consuming, and not scalable for mapping large interaction networks. Therefore, developing efficient computational methods to predict host-pathogen PPIs (HPI) is of great importance.

Currently, many computational approaches have been applied to predict intra-species PPIs [11–14]. However, methods specifically designed for host-pathogen PPIs remain limited. Homology-based methods, such as BIANA, infer new interactions from known PPI data, but struggle to identify additional interactions due to their reliance on existing data [15]. Machine learning models that leverage protein sequence information have improved predictive performance to some extent but are limited by poor generalizability and lack of biological interpretability [16–18]. Recently, deep learning models such as deepHPI have made breakthroughs in HPI prediction by enhancing feature extraction using convolutional neural networks (CNNs) [19]. However, these methods often fail to capture global interaction features and provide limited biological insights. Moreover, most existing models are tailored for single-pathogen systems and do not address the limited coverage of effector-host PPIs in current datasets, which hampers the accurate prediction of bacterial effector-host interactions.

To address these challenges, we developed HPInet, a transformer-based deep learning model specifically designed for predicting bacterial effector-host PPIs. HPInet incorporates convolutional layers into the transformer architecture to enhance local feature extraction and employs attention mechanisms to capture global sequence dependencies. These innovations enable accurate and efficient prediction of effector-host interactions. By leveraging a manually curated dataset of experimentally validated effector-host PPIs, HPInet addresses the limitations of existing models and provides a robust tool for studying cross-species host-pathogen interactions.

In summary, HPInet offers a powerful and interpretable framework for predicting bacterial effector-host interactions, bridging critical gaps in current HPI research. By providing new insights into bacterial pathogenic mechanisms and identifying potential therapeutic targets, HPInet represents a significant advance in the field. The tool is freely available as a web server at https://bis.zju.edu.cn/hpinet.

## Materials and methods

### Data sets collection

A total of 1,855 experimentally validated protein-protein interactions (PPIs) between human proteins and bacterial effector proteins (T1SE-T10SE) were collected from publicly available databases, including HPIDB [20, 21], MINT [22], IntAct [23], PHISTO [24], and relevant literature. After removing duplicates, 1,792 unique PPIs were retained (Supplementary Table S1). To further eliminate redundancy, protein sequences were clustered using CD-HIT [25] with a 40% similarity threshold. For any PPI pair (A-B) resulting in two pairs (A-d) and (c-B) within the same CD-HIT cluster, the pair (c-d) was deemed redundant and excluded. This process yielded a final dataset of 1,552 non-redundant PPIs distributed across T1SEs to T10SEs (Supplementary Figure S1).

For the negative dataset, 10,059 human-bacteria interactions were collected from HPIDB, BV-BRC, IntAct, and PHISTO. Using a “dissimilarity-based negative sampling” strategy [26, 27], bacterial effector proteins were randomly paired with human proteins from the collected interactions. Non-PPIs with sequence similarity exceeding 30% to any known PPI were removed [25]. The final dataset maintained a positive-to-negative ratio of 1:10. The entire dataset was split into training and independent test sets at a 9:1 ratio, ensuring balanced distributions of positive and negative interactions across both sets (Supplementary Figure S1).

### Input embeddings

We utilized two categories of embeddings for protein sequences: frozen embeddings extracted directly from protein language models (pLMs) without fine-tuning on the training dataset and three commonly used protein feature embeddings, namely PSSM, Blosum62, and One-hot. The three pLM-derived embeddings were as follows: **(i) “ESM-2”** [28]: A transformer model with 650 million parameters, pre-trained on UniRef50 [PMID: 25398609]. ESM-2 has demonstrated state-of-the-art performance in protein sequence representation tasks. **(ii) “ProtBert”** [29]: An encoder model based on BERT architecture [30], pre-trained on the UniRef100 dataset [31]. ProtBert leverages the deep bidirectional transformers of BERT to capture contextual features of protein sequences. **(iii) ProtT5-XL** [29]: An encoder model based on the T5 architecture [32], pre-trained on UniRef50. ProtT5-XL extends the capabilities of pLMs for protein sequence representation through its advanced transformer architecture.

### Optimization strategy

To optimize the model’s learning process, we employed the AdamW optimizer combined with a cosine warm-up scheduling strategy, similar to that used in BERT [30]. The initial learning rate was set to 6×10^−5^, with 10 warm-up steps to stabilize the training process.

To address the class imbalance between positive and negative samples, as well as the difficulty of training on hard-to-learn samples, we incorporated Focal Loss [33] into the optimization strategy. Focal Loss (FL) builds upon the weighted Cross-Entropy Loss by introducing a modulating factor controlled by the hyperparameter γ (default γ=2), which adjusts the focus on hard-to-classify samples. This approach ensures more stable gradient descent and improves the model’s ability to handle challenging samples during training.

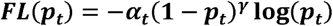

*a_t_*: weight of sample t,

*p_t_*: binary cross entropy loss,

### HPInet model

The HPInet classifier is a deep learning model based on a siamese network architecture, designed for predicting protein-protein interactions (PPIs). The core of the model is a transformer module that integrates convolutional feature extraction and global response normalization (GRN), multi-head self-attention mechanisms to perform protein feature representation learning and classify interaction relationships (Figure 1C). The model takes two sets of protein feature embeddings as input: 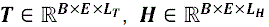, where ***T*** and ***H*** represent embeddings of bacterial and human proteins, respectively. ***B*** denotes the batch size, ***E*** is the embedding dimension, ***L_T_*** and ***L_H_*** are the sequence lengths of bacterial and human proteins.

**Figure 1.**
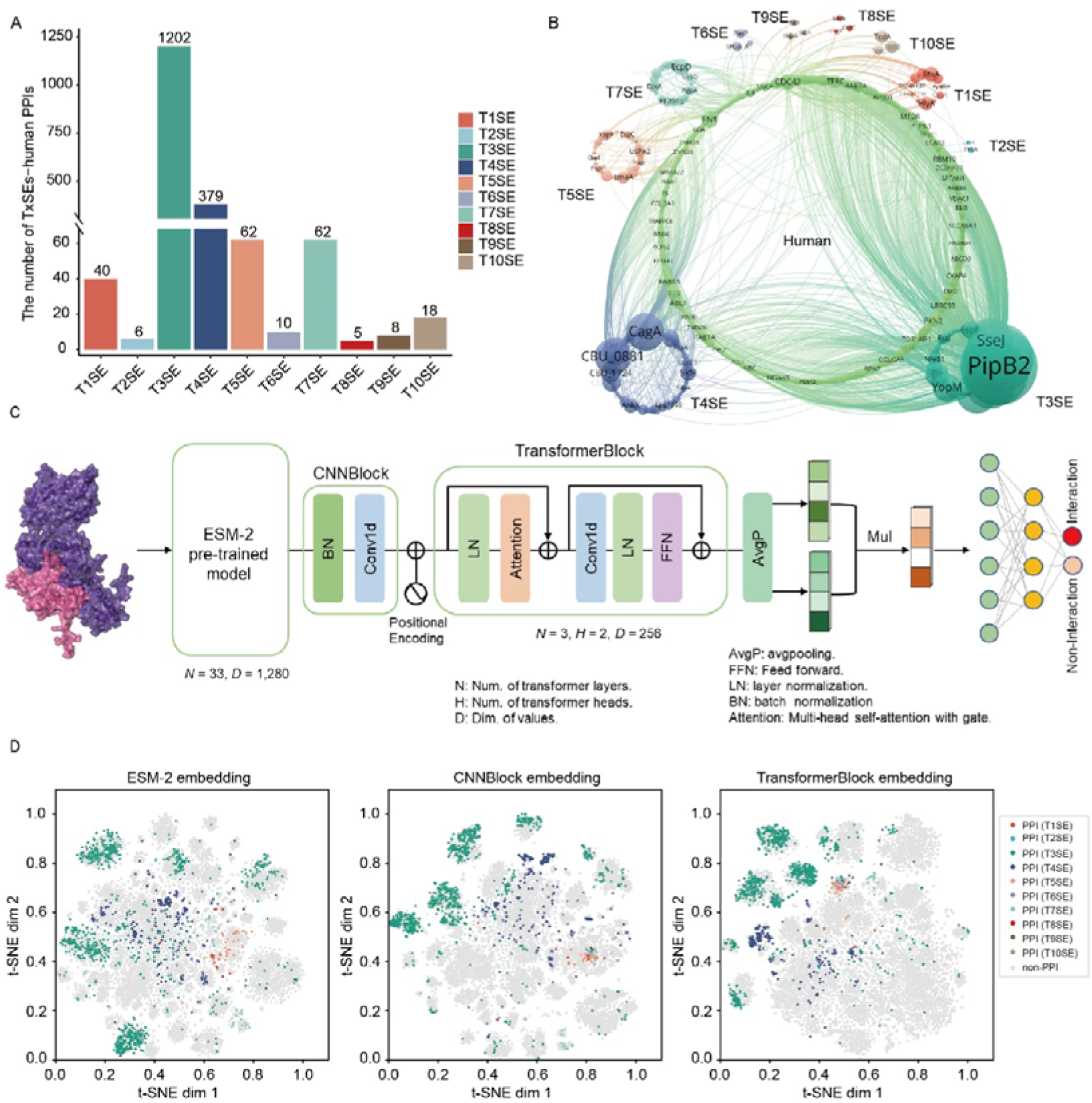
Distribution of human-bacteria secreted protein interactions and HPInet architecture. (A) Distribution of interaction counts between human proteins and bacterial effector proteins from the type I to type X secretion systems. (B) Interaction network of human proteins and bacterial effector proteins. Edges represent protein-protein interactions (PPIs), and node size indicates the number of interactions associated with each protein. (C) Architecture of the HPInet model. Protein sequences are embedded using the ESM-2 model, followed by a convolutional module for extracting local features. A transformer module variant is employed to capture both global and local dependencies, using a gating mechanism to refine interaction predictions. (D) t-SNE visualization of feature embeddings generated at different stages of the model: ESM-2 embeddings (left), embeddings after the convolutional module (middle), and embeddings from the transformer module (right).

Each protein embedding is processed by a shared-weight feature extraction module. Initially, a **CNNBlock** that is 1×1 convolution layer adjusts the feature dimensions:

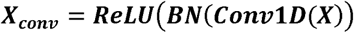

where BN represents batch normalization.

A learnable positional encoding 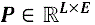 is then added to incorporate sequence information:

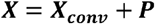

The transformer module subsequently processes ***X*** to extract global dependencies. The transformer employs a gated self-attention mechanism defined as:

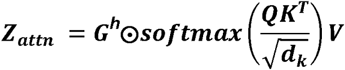

where 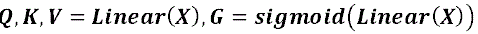, **h** denotes the number of attention heads.

To capture local features, the transformer module incorporates depthwise separable convolution and global response normalization (GRN), inspired by ConvNeXt V2 architecture [34]. These operations are integrated within the feed-forward network (FFN).

The depthwise separable convolution is defined as:

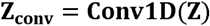

GRN is defined as:

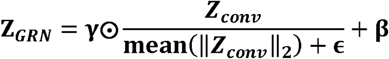

where γ and β are trainable parameters.

To aggregate sequence features into fixed-length representations, mean pooling is applied over the sequence dimension:

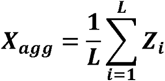

The aggregated representations ***T****_agg_* and ***H****_agg_* are fused via element-wise multiplication:

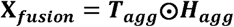

Finally, the fused features, **x***_fusion_* are passed through a feed-forward network (FFN) to predict the interaction probability between the two proteins.

### Performance Evaluation

To evaluate the predictive performance of HPInet and compare it with other models, we employed multiple metrics, including accuracy, sensitivity, specificity, precision, F1-score, Matthews correlation coefficient (MCC), area under the receiver operating characteristic curve (AUROC), and area under the precision-recall curve (AUPRC).

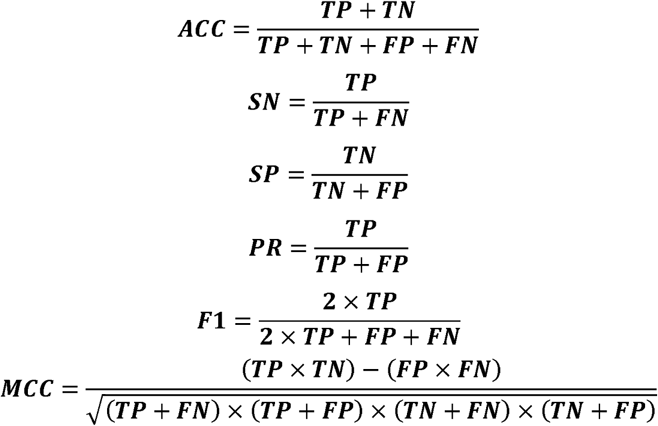

In these metrics, true positives (TP), true negatives (TN), false positives (FP), and false negatives (FN) represent the number of correctly and incorrectly classified samples in the respective categories.

## Results

### 1. Construction of the Human-Bacterial Secreted Protein Interaction Network and Feature Learning with HPInet

Gram-negative bacteria utilize T1SS-T10SS to deliver effector proteins that interact with host proteins, forming a complex network critical for host-pathogen interactions (HPIs) [3–5]. In this study, we collected 1,855 experimentally validated interactions between human proteins and bacterial secreted proteins from multiple databases and literature [20–24]. After redundancy removal, a final dataset of 1,552 non-redundant interactions was obtained (Figure 1A). These interactions spanned all secretion systems effectors (T1SEs-T10SEs), with the majority originating from T3SE and T4SE, accounting for 1,202 and 379 interactions, respectively (Figure 1B). This indicates the prominent roles of type III and type IV secretion systems in bacterial infections and highlights the diversity in interaction patterns mediated by different secretion systems. For example, critical effector proteins such as CagA (type IV secretion effector) and PipB2 (type III secretion effector) interact with multiple host target proteins, reflecting their importance in pathogenic mechanisms.

To further investigate the interaction patterns and features of these effector-host protein pairs, we developed HPInet, a deep learning model based on a siamese network architecture. HPInet is specifically designed to predict interactions between bacterial effector proteins and host proteins. The model integrates convolutional layers and a variant of the transformer module to enable hierarchical feature learning, capturing both local and global dependencies (Figure 1C).

Initially, protein sequences were embedded into high-dimensional feature vectors using the pre-trained ESM-2 model [PMID: 36927031], capturing fundamental sequence features. Next, a convolutional module extracted local feature patterns from these embeddings. The transformer module, enhanced with multi-head self-attention and depthwise separable convolutions and GRN in its feed-forward network (FFN), simultaneously captured both global and local dependencies. The module also employed a gating mechanism to selectively weight features. Finally, the model fused features from the two proteins and outputted classification probabilities for PPI prediction. This design enabled HPInet to efficiently capture the complex interaction patterns between proteins while enhancing model generalization and interpretability.

To validate the feature learning capability of HPInet, we performed t-SNE visualizations of the embeddings at different stages of the model (Figure 1D). The initial embeddings generated by ESM-2 (left) reflected basic sequence features but failed to clearly separate PPIs from non-PPIs. After processing by the convolutional module (middle), feature separation improved. Finally, embeddings derived from the transformer module (right) demonstrated distinct clustering in the feature space, with clear separation of interactions across T1SEs-T10SEs. These results indicate that HPInet progressively refines feature representations, evolving from basic sequence information to a nuanced understanding of complex interaction patterns, achieving precise predictions for bacterial effector-host protein interactions.

### 2. Transformer-Based Deep Learning Prediction Model: HPInet

Recent advances in protein language models have revolutionized biological sequence classification and structural prediction tasks. We evaluated HPInet using six embedding approaches, including three pretrained protein language models (ESM-2, ProtBert, ProtT5-XL-U50) [28, 29] and three traditional embeddings (PSSM, Blosum62, and One-hot). These features were systematically evaluated using the HPInet model, which is based on a transformer architecture.

Among all embeddings tested, HPInet (ESM-2) using ESM-2 embeddings consistently achieved the best performance across validation and independent test sets, with the highest accuracy, specificity, precision, F1-score, MCC, AUROC, and AUPRC (Figures 2A-B, Supplementary Table S2 and S3). In the independent test set, HPInet (ESM-2) achieved an average accuracy of 0.882, an F1-score of 0.532, and an AUPRC of 0.522, significantly outperforming models trained on other feature embeddings.

**Figure 2.**
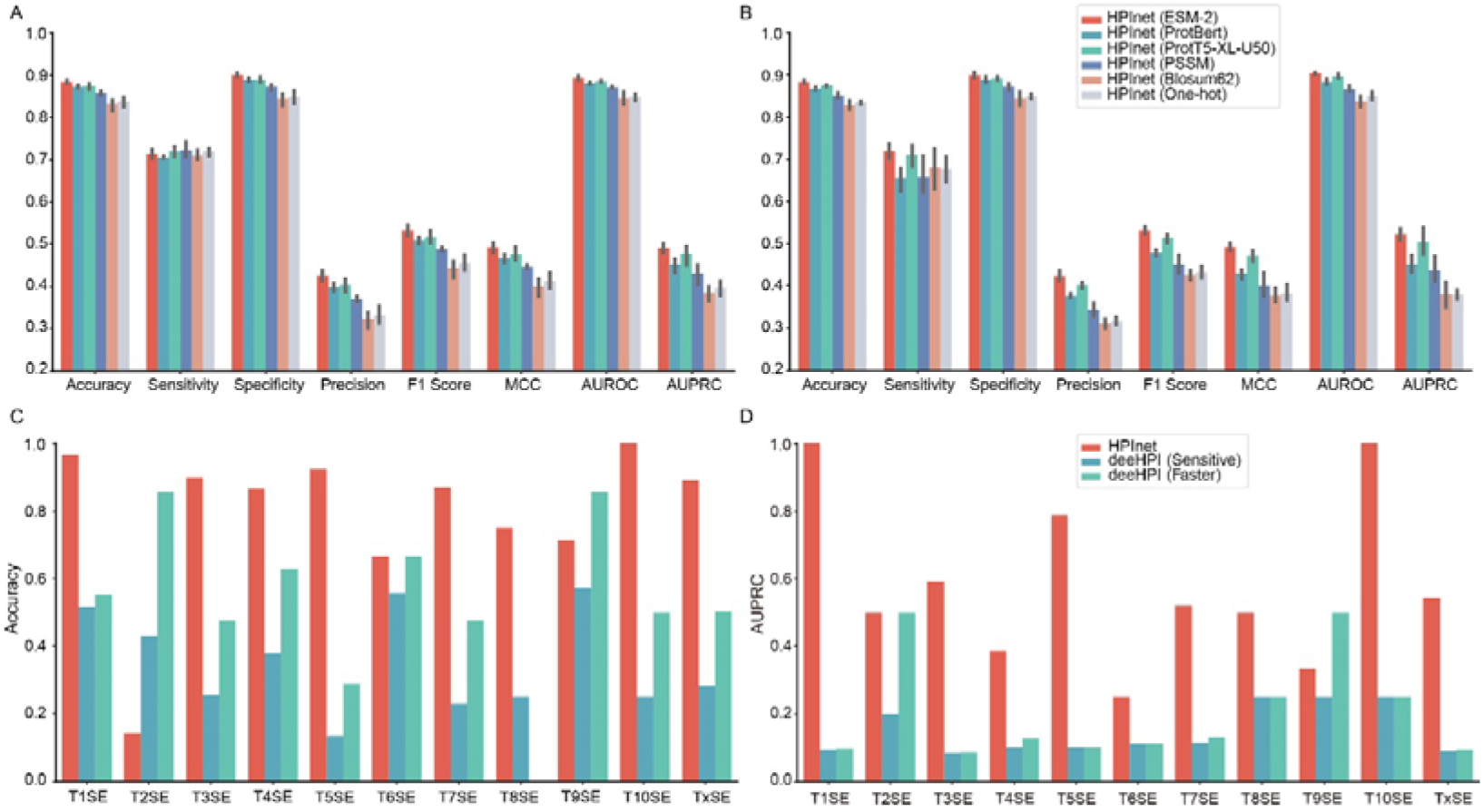
Performance evaluation of HPInet and comparison with deepHPI across different types of secreted proteins. (A) Performance of HPInet models trained with six different protein feature embeddings, evaluated on the validation set using metrics including Accuracy, Sensitivity, Specificity, Precision, F1-score, MCC, AUROC, and AUPRC. (B) Performance of HPInet models on the independent test set. (C-D) Comparative evaluation of HPInet (ESM-2) and deepHPI (Sensitive and Faster modes) across 10 types of secreted proteins (T1SE-T10SE) on the independent test set, focusing on (C) Accuracy and (D) AUPRC.

To further validate the performance of HPInet, we compared it against leading host-pathogen PPI prediction tools, including BIANA [15], Mei’s method [16], Ahmed’s method [17], InterSPPI [18], and deepHPI [19]. While most of these tools are tailored to predict PPIs for specific pathogens, only deepHPI supports cross-species prediction [19]. Therefore, we primarily compared HPInet with the two modes of deepHPI (Sensitive and Faster) (Figures 2C-D, Supplementary Table S4).

The results showed that deepHPI (Faster) outperformed deepHPI (Sensitive) in overall performance, so we used deepHPI (Faster) as the primary comparator. On the independent test set, HPInet-ESM-2 significantly outperformed deepHPI (Faster), with accuracy improving from 0.502 to 0.891, F1-score from 0.161 to 0.543, and AUPRC from 0.095 to 0.522 (Figures 2C-D, Supplementary Table S4). Furthermore, we evaluated the performance of HPInet and deepHPI across T1SEs to T10SEs. Except for T2SE, HPInet outperformed deepHPI (Faster) across all secretion system types, with particularly notable improvements in T1SE, T3SE, and T10SE predictions.

These findings demonstrate that HPInet effectively captures complex interaction features between diverse bacterial effector proteins and host proteins, providing a significant advancement in PPI prediction accuracy and reliability.

### 3. Ablation Experiments to Validate the Effectiveness of HPInet Key Modules

To evaluate the contributions of key modules in HPInet, we conducted ablation experiments comparing the full HPInet model with two variant models: **HPInet (basic Transformer, baseline)**, which excludes CNN and GRN from the transformer’s FFN, and **HPInet (without Transformer, w/o)**, which removes the transformer module entirely and relies solely on convolutional layers for feature extraction. The results from 5-fold cross-validation (CV) and independent test sets (Ind. Tests) are presented in Table 1.

**Table 1.**
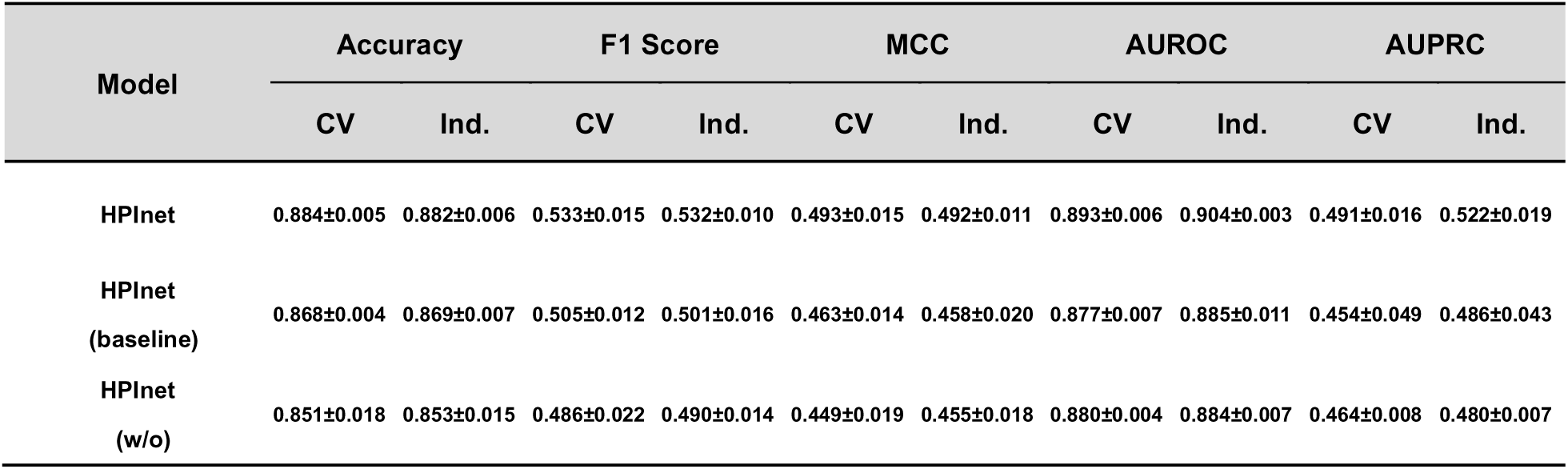
Performance of ablation experiments on different model architectures in 5-fold cross-validation (CV) and independent (Ind.) Tests.

| Model | Accuracy |  | F1 Score |  | MCC |  | AUROC |  | AUPRC |  |
| --- | --- | --- | --- | --- | --- | --- | --- | --- | --- | --- |
|  | CV | Ind. | CV | Ind. | CV | Ind. | CV | Ind. | CV | Ind. |
| HPINet | 0.884±0.005 | 0.882±0.006 | 0.533±0.015 | 0.532±0.010 | 0.493±0.015 | 0.492±0.011 | 0.893±0.006 | 0.904±0.003 | 0.491±0.016 | 0.522±0.019 |
| HPINet<br>(baseline) | 0.868±0.004 | 0.869±0.007 | 0.505±0.012 | 0.501±0.016 | 0.463±0.014 | 0.458±0.020 | 0.877±0.007 | 0.885±0.011 | 0.454±0.049 | 0.486±0.043 |
| HPINet<br>(w/o) | 0.851±0.018 | 0.853±0.015 | 0.486±0.022 | 0.490±0.014 | 0.449±0.019 | 0.455±0.018 | 0.880±0.004 | 0.884±0.007 | 0.464±0.008 | 0.480±0.007 |

The experimental results demonstrate that the complete HPInet model outperformed both variants across all evaluation metrics. On the independent test set, HPInet achieved an AUPRC of 0.522, significantly higher than HPInet (baseline) at 0.478 and HPInet (w/o) at 0.431. These findings indicate that integrating CNN and GRN modules into the transformer’s FFN enhances the model’s ability to extract local features and normalize global features effectively. Moreover, the inclusion of the transformer module is critical for capturing the complex dependencies between interacting proteins. In contrast, models without the transformer showed a substantial performance decline, further underscoring the importance of the transformer module.

In summary, the ablation experiments validate the effectiveness of HPInet’s architecture. The combination of the transformer module with CNN and GRN modules significantly contributes to improving the accuracy and generalizability of predictions for bacterial effector-host protein interactions.

### 4. Interpretability of HPInet and Comparison with Docking Tools

To further assess the interpretability of HPInet, we performed docking analyses using AlphaFold3 [35] on interactions between bacterial effector proteins and human host proteins. For each PPI, we selected the top-ranked docking structure (Top1) from AlphaFold3 predictions and filtered those with a predicted Template Modeling (pTM) score > 0.5, considering these as reliable global structural predictions. Using HPInet, we identified key interaction sites based on self-attention scores, defining residues with attention scores above the median as significant interaction residues. Additionally, SHAP’s GradientShap model was applied to further interpret key residues [36]. Binding sites predicted by PDBePISA (https://www.ebi.ac.uk/pdbe/pisa/) from AlphaFold3 docking results were used as the ground truth. For each PPI, we calculated precision values for key residues and analyzed their correlation with AlphaFold3’s ipTM scores (Figure 3A). The results revealed a significant positive correlation between the precision values of HPInet’s predicted key residues and AlphaFold3’s ipTM scores (GradientShap *R*=0.35, *P*=2.94e-20; Attention *R*=0.22, *P*=5.80e-09), indicating that HPInet effectively captures important interaction sites consistent with AlphaFold3.

**Figure 3.**
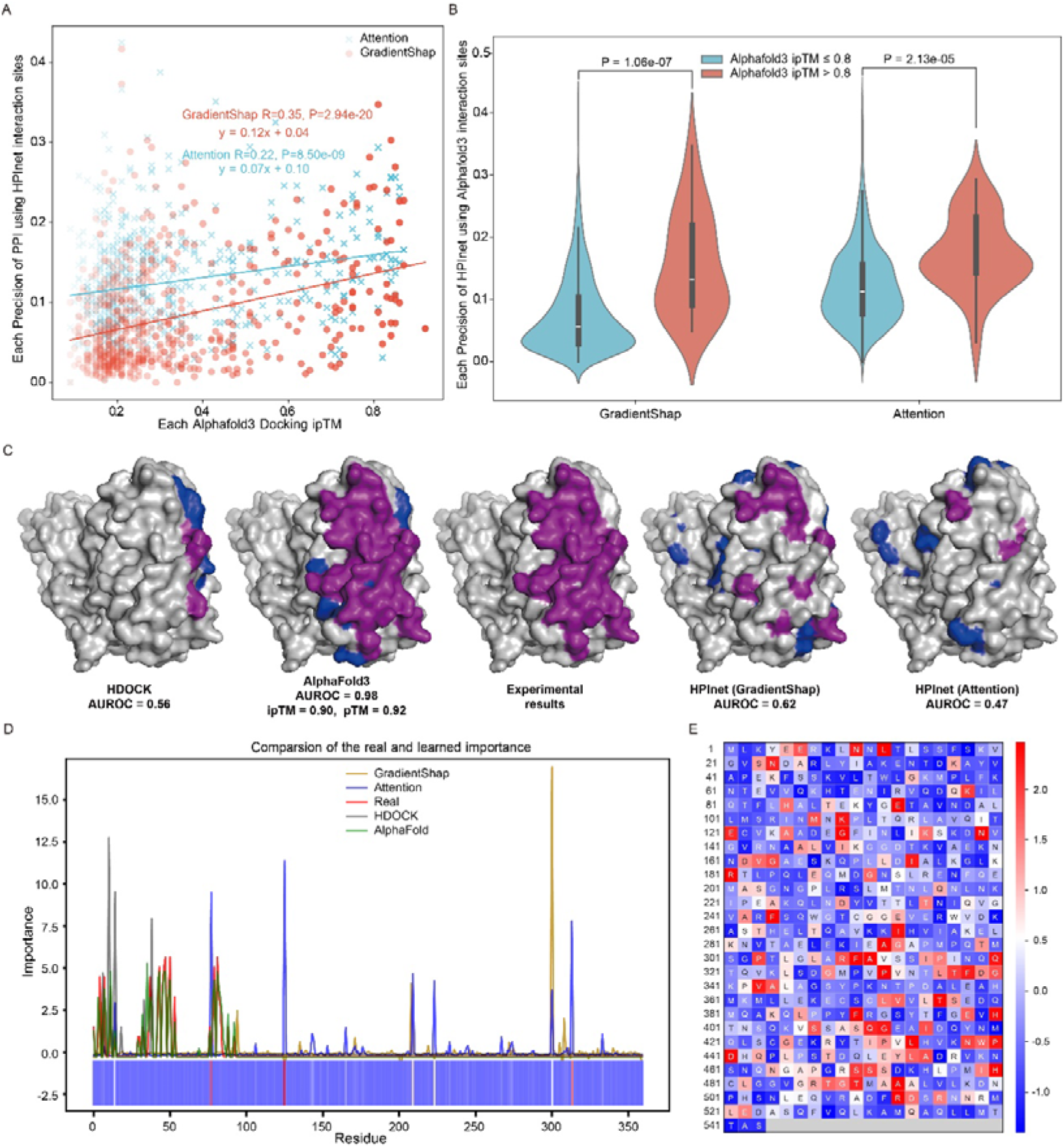
Comparison of HPInet and docking tools in residue-level interaction prediction. (A) Correlation analysis between the precision of important interaction residues predicted by HPInet (using GradientShap and attention scores) and AlphaFold3’s ipTM values. (B) Distribution of HPInet predictions (GradientShap and attention scores) grouped by AlphaFold3 ipTM values, with 0.8 used as the threshold. (C) Comparative performance of different tools, including AlphaFold3, HDock, HPInet (GradientShap), and HPInet (Attention), in predicting interaction residues for the SptP-Rac1 complex (PDB: 1g4u-S). AUROC values, calculated using residues identified by PDBePISA as ground truth, are used to evaluate predictive performance. Pink and blue indicate residues that are the same as or different from the interaction sites predicted by the crystal structure, respectively. (D) Distribution of interaction residues in the crystal structure of SptP (PDB: 1g4u-S). “Real” indicates important residues identified by PDBePISA as critical interaction sites, normalized using Z-scores. The heatmap represents interaction residues inferred byHPInet using attention scores. (E) Heatmap of attention-derived important residues predicted by HPInet for the full-length SptP protein in the SptP-Rac1 interaction.

Notably, although AlphaFold3 considers predictions with pTM > 0.5 and ipTM > 0.8 to reflect reliable docking outcomes, our analysis of all 1,792 PPIs predicted by AlphaFold3 identified only 17 pairs meeting these criteria (Supplementary Table S5). Strikingly, 7 of these 17 pairs (41.2%) corresponded to experimentally validated crystal structures in the RCSB PDB database (https://www.rcsb.org/, Supplementary Table S5). This highlights the inherent limitations of structure-based docking approaches like AlphaFold3 in predicting interactions between bacterial secretory proteins and host proteins, as well as in identifying their interaction sites.

Further comparisons of PPIs with high (ipTM>0.8) and low (ipTM≤0.8) scores (Figure 3B) demonstrated that HPInet provided more stable predictions for PPIs with high ipTM scores (*P*<0.05), suggesting that the model exhibits stronger interpretability under reliable docking conditions.

We also retrieved a crystal structure of an interaction pair from the RCSB PDB database (PDB ID: 1g4u), representing *Salmonella* effector protein SptP (P74873) and human GTPase Rac1 (P63000) [37]. In the 1g4u structure, SptP contains two residues 167-461 and 474-539. We compared the predictive performance of AlphaFold3, HDOCK [38], and HPInet in identifying key interaction sites. AlphaFold3 achieved the highest AUROC (0.98), followed by HPInet using GradientShap (AUROC = 0.62), which outperformed HDOCK (AUROC = 0.56) (Figure 3C). Although HPInet’s AUROC based on attention scores was slightly lower (0.47), the model’s multi-dimensional interpretability provides valuable insights into PPI mechanisms (Figure 3C).

We further analyzed the overlap between the key interaction sites predicted by AlphaFold3, HDOCK, and HPInet and those identified in the 1g4u crystal structure (Figure 3D). HPInet’s attention-based predictions showed high consistency (AUROC = 0.50) with AlphaFold3 for residues within the first 123 amino acids (167-290) of SptP, a region corresponding to its GTPase-activating protein (GAP) activity. This region has been experimentally validated as a critical interaction site [37, 39]. However, the overlap decreased for residues in the tyrosine phosphatase domain of SptP, which plays a crucial role during bacterial infection by targeting tyrosine-phosphorylated host proteins. This domain is also believed to target host substrates such as Cdc42 [37]. Notably, the phosphatase active site of SptP is located at residues 479-489, with Cys-481 serving as the catalytic residue [37]. This active site coincided with HPInet’s attention-derived key residues (Figure 3E).

These findings demonstrate that HPInet, while not surpassing AlphaFold3 in structural accuracy, provides complementary and biologically meaningful insights into interaction sites, especially in regions critical for effector-host interactions.

### 5. Performance of HPInet in predicting interactions between different bacterial effectors and human host proteins

We systematically analyzed the performance of HPInet in predicting interactions between different bacterial effector proteins and human host proteins, focusing on comparisons between models with and without fine-tuning (Table 2). Due to the limited number of interactions available for certain secretion systems (Figure 1A), we trained separate models for T3SE, T4SE, T5SE, and T7SE interactions with host proteins. The results showed that fine-tuning consistently improved the predictive performance across all secretion systems, particularly in terms of AUPRC. However, the extent of improvement varied depending on the secretion system (Table 2).

**Table 2.** Performance Before and After Model Finetuning (FT) in 5-Fold Cross-Validation (CV) and Independent (Ind.) Tests.

**Cross-Validation (CV) and Independent (Ind.) Tests.**
| Model<br>(HPINet) | Accuracy |  | F1 Score |  | MCC |  | AUROC |  | AUPRC |  |
| --- | --- | --- | --- | --- | --- | --- | --- | --- | --- | --- |
|  | CV | Ind. | CV | Ind. | CV | Ind. | CV | Ind. | CV | Ind. |
| T3SE | 0.862±0.017 | 0.873±0.015 | 0.505±0.014 | 0.549±0.014 | 0.476±0.006 | **0.529±0.010** | **0.898±0.011** | **0.926±0.004** | **0.478±0.014** | 0.565±0.021 |
| T3SE (FT) | **0.879±0.008** | **0.885±0.008** | **0.525±0.009** | **0.554±0.014** | **0.489±0.008** | 0.523±0.013 | 0.897±0.011 | 0.923±0.004 | 0.473±0.017 | **0.566±0.017** |
| T4SE | 0.797±0.031 | 0.780±0.031 | 0.388±0.023 | 0.381±0.021 | 0.344±0.019 | 0.339±0.020 | 0.813±0.014 | 0.814±0.016 | 0.388±0.059 | 0.328±0.036 |
| T4SE (FT) | **0.842±0.044** | **0.818±0.033** | **0.461±0.067** | **0.405±0.021** | **0.422±0.067** | **0.357±0.019** | **0.842±0.030** | **0.824±0.016** | **0.442±0.088** | **0.349±0.019** |
| T5SE | **0.915±0.039** | **0.912±0.011** | 0.623±0.163 | **0.622±0.050** | 0.593±0.178 | **0.587±0.058** | 0.893±0.088 | 0.918±0.019 | **0.711±0.156** | 0.651±0.136 |
| T5SE (FT) | 0.911±0.064 | 0.900±0.044 | **0.643±0.179** | 0.603±0.079 | **0.616±0.185** | 0.575±0.077 | **0.900±0.049** | **0.936±0.017** | 0.672±0.159 | **0.713±0.090** |
| T7SE | 0.837±0.051 | 0.836±0.052 | 0.427±0.083 | 0.369±0.145 | 0.381±0.104 | 0.291±0.166 | 0.819±0.047 | 0.724±0.059 | 0.451±0.053 | 0.390±0.138 |
| T7SE (FT) | **0.874±0.019** | **0.885±0.058** | **0.506±0.074** | **0.503±0.135** | **0.468±0.093** | **0.459±0.172** | **0.866±0.051** | **0.817±0.097** | **0.524±0.145** | **0.548±0.165** |

For example, in the independent test set, fine-tuning resulted in relatively small AUPRC improvements for HPInet (T3SE) and HPInet (T4SE), with increases of 0.18% and 6.40%, respectively. This limited improvement may be attributed to the predominance of T3SE and T4SE interactions in the training dataset, which allowed the models to achieve high feature adaptation even before fine-tuning. In contrast, fine-tuning led to substantial improvements for HPInet (T5SE) and HPInet (T7SE), with AUPRC increases of 9.52% and 40.51%, respectively. This suggests that fine-tuning not only enhances the model’s ability to learn data-specific features but also optimizes the detection of shared interaction patterns between T5SE and T7SE effectors and host proteins.

We further evaluated HPInet’s generalization ability in cross-secretion system predictions (Figures 4A-B). Predictions within the same secretion system consistently outperformed cross-system predictions. However, the cross-system performance between T5SE and T7SE was significantly better than other secretion system combinations (Figure 4A). Without fine-tuning, the AUPRC for the T5SE model predicting T7SE interactions was 0.60, and for the T7SE model predicting T5SE interactions, it was 0.64. After fine-tuning, the AUPRC for both scenarios improved to 0.71 (Figure 4B). These results indicate that T5SE and T7SE likely share critical features, enabling more efficient cross-system transfer learning.

**Figure 4.**
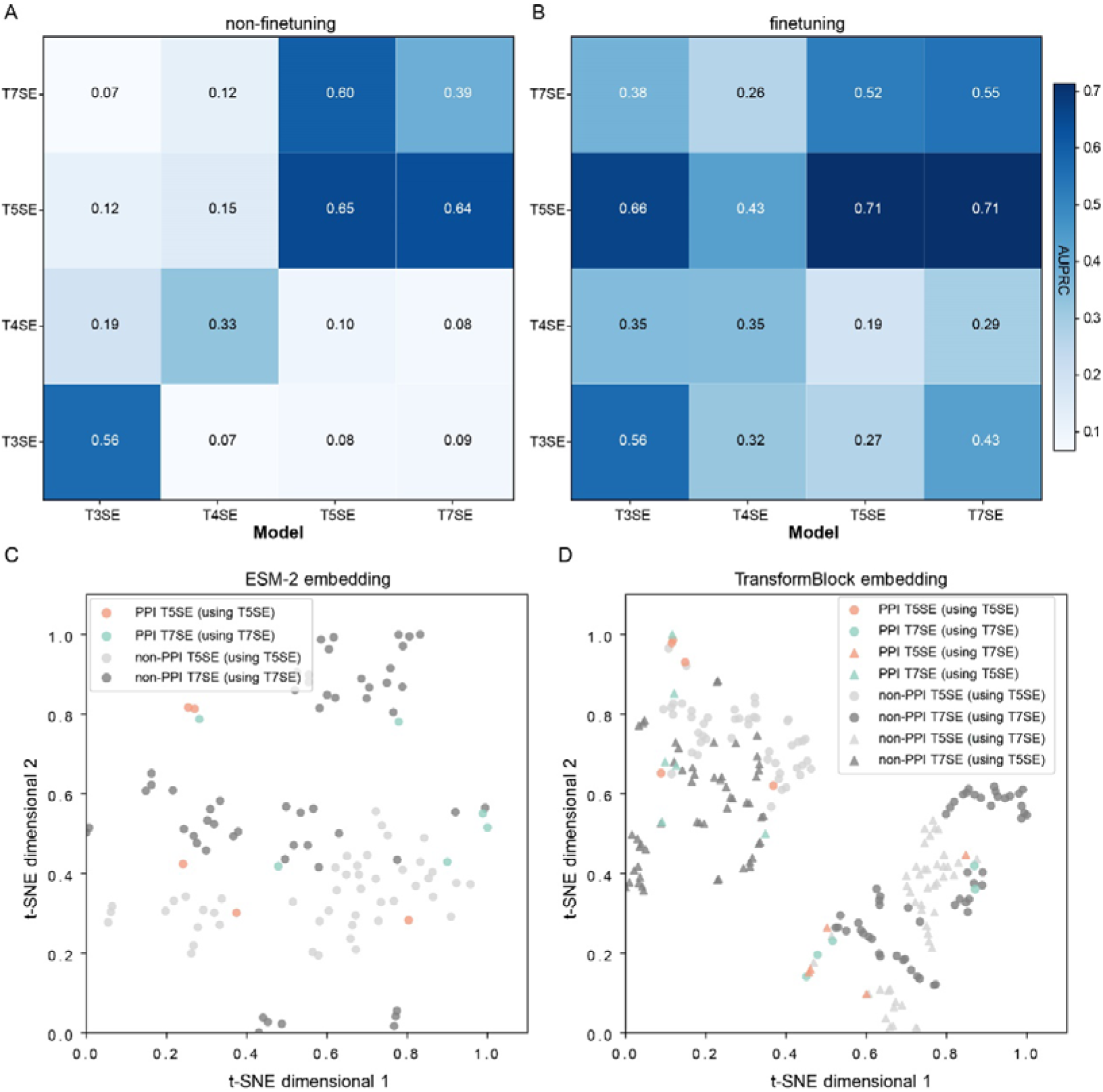
Analysis of HPInet prediction performance and feature space for HPI across different secretion systems. (A-B) AUPRC performance of HPInet in cross-secretion system predictions without fine-tuning (A) and with fine-tuning (B). (C-D) t-SNE clustering of T5SE and T7SE interaction data at the input layer (ESM-2 embeddings) (C) and at the output of the transformer module (D). The shape of each point indicates whether the same (circle) or different (triangle) secretion protein-host interaction data were used in training versus testing.

To explore the superior cross-system performance between T5SE and T7SE, we conducted t-SNE clustering analysis on features from the input layer (ESM-2 embeddings) and the output layer of the transformer module (Figures 4C-D). The input feature space (Figure 4C) revealed significant differences in the feature distributions of T5SE and T7SE interactions, suggesting initial sequence-level disparities. However, in the transformer module’s output feature space (Figure 4D), a single model exhibited strong similarity in its representation of T5SE and T7SE interactions, while distinct models showed notable differences in feature distributions. This indicates that HPInet effectively captures shared features between secretion systems while retaining system-specific characteristics in its output layer.

These findings demonstrate HPInet’s ability to generalize across secretion systems, particularly between T5SE and T7SE, where the model benefits from fine-tuning to enhance its learning of shared patterns and improve predictive performance.

### 6. HPInet web server and implementation

To ensure the accessibility and usability of HPInet, we developed a web-based application that provides a user-friendly interface for protein-protein interaction (PPI) prediction (Figure 5). The web server backend is implemented in PHP, with the frontend designed using HTML and JavaScript.

**Figure 5.**
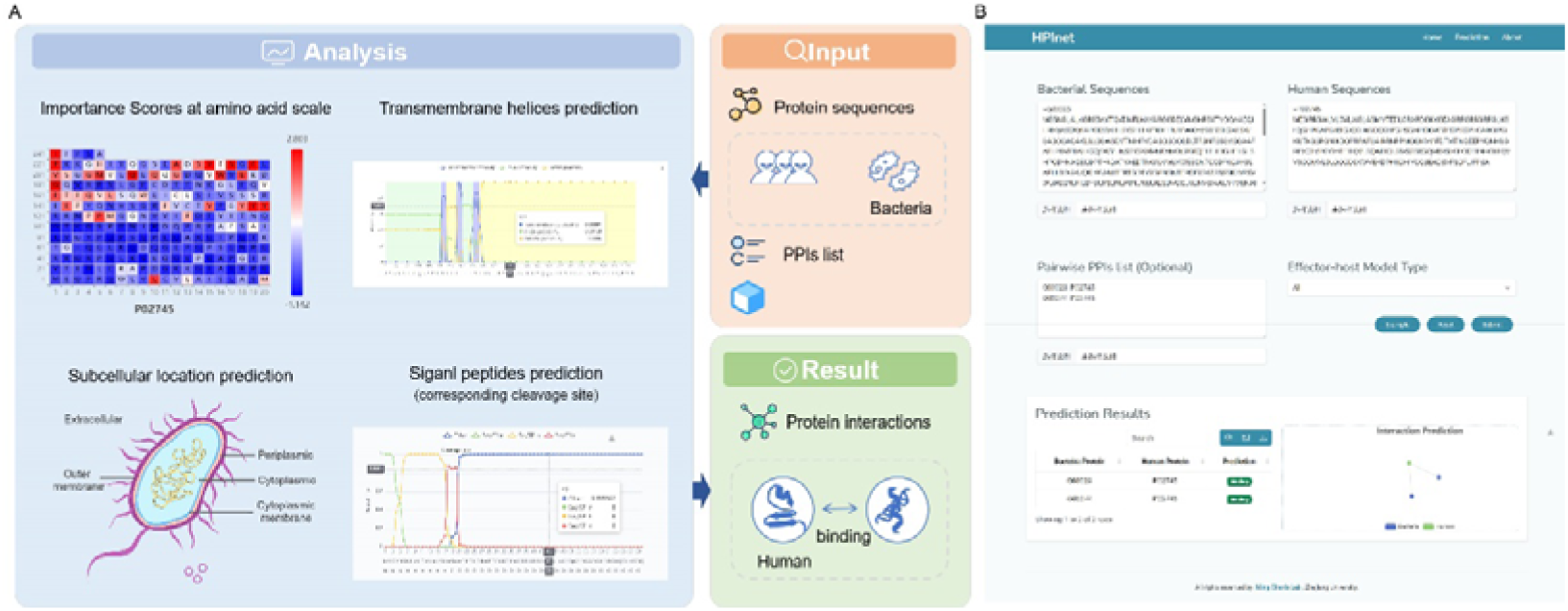
Overview of HPInet web server for human-bacteria protein-protein interaction (PPI) prediction. (A)The HPInet framework comprises three main sections: Input, Analysis, and Result. (B) The prediction server of HPInet, showing input fields for bacterial and human protein sequences, optional PPI lists, and effector-host model selection. The output includes predicted binding interactions displayed in tabular format and as an interaction network visualization.

Users can submit one or multiple protein sequences in FASTA format for both bacterial effector proteins and host (human) proteins. The server offers a selection of models, including HPInet, HPInet(T3SE), HPInet(T4SE), HPInet(T5SE), and HPInet(T7SE), to meet specific prediction needs (Figure 5B). Prediction results are available for download in CSV format, with sortable columns and keyword-based filtering for efficient result exploration (Figure 5B). Additionally, an interactive PPI network is visualized using Echarts. To provide users with deeper insights, they can click on any row in the results table to view detailed interaction information. The detailed results page includes the following scores for each protein pair, with separate data for bacterial and human proteins: importance scores, transmembrane helices, signal peptides, and subcellular location (Figure 5A).

This web application provides an intuitive and efficient platform for researchers to explore host-pathogen protein interactions and is freely available at https://bis.zju.edu.cn/hpinet.

## Discussion

Bacterial effector proteins secreted through various secretion systems (T1SS-T10SS) are critical to host-pathogen interactions, manipulating host cellular processes to facilitate infection and disease [3–5]. Understanding these interactions is essential for uncovering bacterial pathogenic mechanisms and identifying potential therapeutic targets. Despite their importance, experimental methods for identifying effector-host protein-protein interactions (PPIs) are time-consuming, labor-intensive, and limited in scale, creating a pressing need for accurate computational approaches. To address this, we developed HPInet, a transformer-based deep learning model designed to predict PPIs between bacterial effector proteins and human host proteins.

HPInet outperformed existing methods, including deepHPI, achieving significant improvements in accuracy (from 0.502 to 0.891) and sensitivity (35.3%). By integrating convolutional layers and GRN into the transformer’s FFN, HPInet effectively captured both local sequence features and global dependencies, enabling high-precision predictions across diverse bacterial secretion systems. The model’s attention mechanism further enhanced interpretability by identifying key residues involved in protein interactions, offering valuable insights for experimental validation.

Our analyses revealed that HPInet performs particularly well in predicting PPIs within individual secretion systems, with notable improvements after fine-tuning. For instance, the AUPRC of T7SE predictions improved by 40.51% following fine-tuning, highlighting the model’s ability to learn and adapt to specific data distributions. Interestingly, HPInet also demonstrated strong cross-system predictive capabilities, particularly between T5SEs and T7SEs, likely due to shared secretion mechanisms. These findings underscore HPInet’s ability to generalize while retaining system-specific features, as evidenced by t-SNE clustering analyses, which showed distinct input feature distributions but convergent output representations for T5SEs and T7SEs.

HPInet’s interpretability was further validated by its ability to predict biologically relevant interaction sites. For example, in the case of the interaction between the *Salmonella* type III secreted effector SptP and human Rac1 interact [37, 39], HPInet identified residues overlapping with experimentally validated binding sites, including the tyrosine-protein phosphatase domain, a critical region for bacterial virulence [37]. This highlights HPInet’s potential for elucidating effector-host interaction mechanisms and prioritizing therapeutic targets.

Despite its advancements, HPInet faces several challenges. The scarcity of high-quality experimental datasets, especially for underrepresented secretion systems, limits its generalizability. Incorporating structural data from tools like AlphaFold3 [35] or expanding to multi-host predictions could further enhance its utility. Additionally, while HPInet effectively captures shared and system-specific features, the biological significance of some model-learned patterns remains to be explored.

In conclusion, HPInet offers a powerful and interpretable framework for host-pathogen PPI prediction, bridging critical gaps in our understanding of bacterial effector-host interactions. Its robust performance, combined with its deployment as a web server, provides a valuable resource for researchers to investigate bacterial pathogenesis and identify novel therapeutic strategies. By addressing current limitations and expanding its scope, HPInet holds significant potential for advancing research in host-pathogen biology.

## Key Points

**Novel Framework with Enhanced Feature Extraction:** HPInet is a novel transformer-based deep learning framework that integrates convolutional layers and global response normalization. This combined approach enables the model to effectively capture both local sequence features and global dependencies, which is critical for accurately predicting protein-protein interactions between bacterial effector proteins and human host proteins.

**Superior Predictive Performance:** The model significantly improves prediction accuracy (from 0.502 to 0.891) and boosts sensitivity by 35.3% compared to existing methods.

**Biological Interpretability:** HPInet’s attention mechanism allows it to identify critical interaction residues, providing insights that align with experimentally validated binding sites.

**Accessible Tool for Research:** Deployed as a free web server, HPInet offers a practical and interpretable resource for researchers studying bacterial pathogenesis and potential therapeutic targets.

## Data availability

### Availability

The online version of HPInet is freely accessible at http://bis.zju.edu.cn/HPInet.

## Authors’ Contribution

MC, and YH conceived and supervised the project. YH, MC, and YW coordinated the project. YH, and YZ dataset collection. YH provided codes, models and software tools. YH, and LL developed the website. YH, LL, YZ, EL, HC, SL, CF, YSH, YC, SZ, LX, YW, and MC wrote the first draft of this manuscript. YH, YW, and MC revised the manuscript accordingly.

## Author Biographies

**Yueming Hu** is a PhD candidate at Department of Bioinformatics, College of Life Sciences, Zhejiang University. He researches bioinformatics with a focus on deep learning and pathogenic bacterial systems biology.

**Liya Liu** is a Master’s candidate at Department of Bioinformatics, College of Life Sciences, Zhejiang University. Her research centers on bioinformatics analysis

**Yanyan Zhu** is a Master’s candidate at Department of Bioinformatics, College of Life Sciences, Zhejiang University. Her work emphasizes bioinformatics and pathogenic bacterial systems biology.

**Enyan Liu** is a PhD candidate at Department of Bioinformatics, College of Life Sciences, Zhejiang University. Her research focuses on the bioinformatics and deep learning.

**Haoyu Chao** is a PhD candidate at Department of Bioinformatics, College of Life Sciences, Zhejiang University. His work centers on the bioinformatics data analysis.

**Sida Li** is a PhD candidate at Department of Bioinformatics, College of Life Sciences, Zhejiang University. His research focuses on the bioinformatics data analysis.

**Cong Feng** is a postdoctoral research fellow at the Department of Bioinformatics, College of Life Sciences, Zhejiang University. His research focuses on DNA repeat detection and genome analysis.

**Yanshi Hu** is a postdoctoral research fellow at the Department of Bioinformatics, College of Life Sciences, Zhejiang University. His research focuses on the bioinformatics data analysis.

**Yuhao Chen** is a PhD candidate at Department of Bioinformatics, College of Life Sciences, Zhejiang University. His research focuses on the bioinformatics and deep learning.

**Shilong Zhang** is a PhD candidate at Department of Bioinformatics, College of Life Sciences, Zhejiang University. His research focuses on the bioinformatics and deep learning.

**Yifan Chen** is a PhD candidate at Department of Bioinformatics, College of Life Sciences, Zhejiang University. Her research focuse on the bioinformatics analysis and deep learning.

**Luyao Xie** is a Master candidate at Department of Bioinformatics, College of Life Sciences, Zhejiang University. Her research centers on the bioinformatics analysis.

**Yejun Wang** is an associate professor in the College of Basic Medicine, Shenzhen University Medical School. His research focuses on bacterial secretion systems, precision oncology, genome assembly, and deep learning for predicting pathogenic proteins and antigens.

**Ming Chen** is a full professor in the Department of Bioinformatics, College of Life Sciences, Zhejiang University. He is the president of Bioinformatics Society of Zhejiang Province, China. His current research focuses on establishing useful bioinformatics tools and platforms to integrate and analyze massive biological datasets.

## Conflict of Interest: none declared

## Supplementary data

**Supplementary Figure S1.**
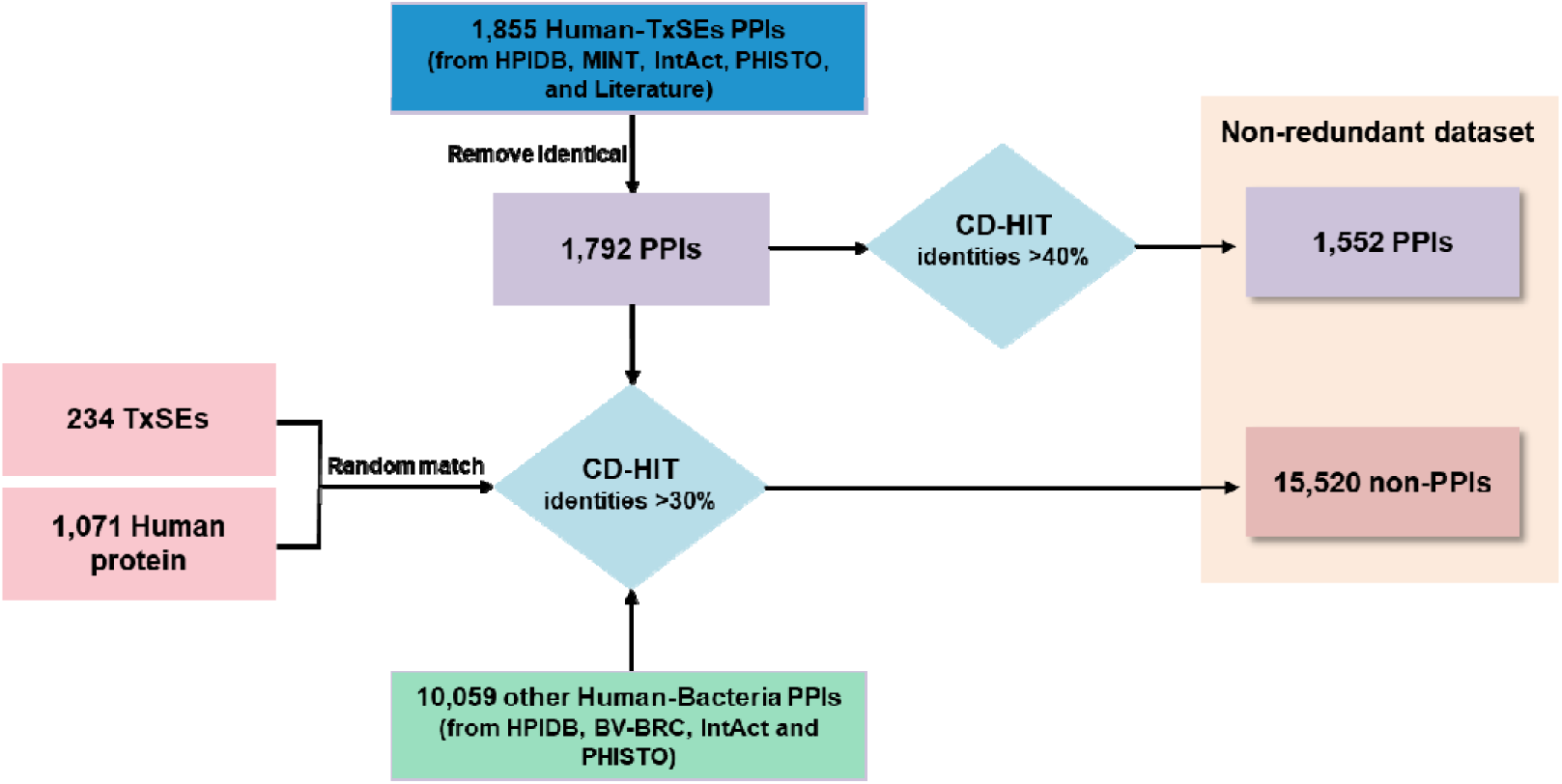
The workflow to construct the training and independent testing dataset in this study.

**Supplementary Table S1.** Experimentally validated protein-protein interactions (PPIs) between bacterial secretion system effector proteins and human proteins.

**Supplementary Table S2.** Performance comparison of the models in HPInet on 5-fold cross-validation dataset.

**Supplementary Table S3.** Performance comparison of the models in HPInet on Independent Testing dataset.

**Supplementary Table S4.** Benchmarking performance between HPInet and two operational modes of deepHPI on on Independent Testing dataset.

**Supplementary Table S5.** Experimentally validated crystal structures in the PDB database among high-confidence AlphaFold3-predicted bacterial effector-human PPIs (pTM >0.5, ipTM >0.8).

